# Gut-associated microbes are present and active in the pig nasal cavity

**DOI:** 10.1101/2023.06.12.544581

**Authors:** Pau Obregon-Gutierrez, Laura Bonillo-Lopez, Florencia Correa-Fiz, Marina Sibila, Joaquim Segalés, Karl Kochanowski, Virginia Aragon

**Author notes:** shared-first authors (Pau Obregon-Gutierrez, Laura Bonillo-Lopez). shared-last authors (,).

## Abstract

**Background:** The nasal microbiota is a key contributor to animal health, and characterizing the nasal microbiota composition is an important step towards elucidating the role of its different members. Efforts to characterize the nasal microbiota composition of domestic pigs and other farm animals frequently report the presence of bacteria that are typically found in the gut, including many anaerobes from the *Bacteroidales* and *Clostridiales* orders. However, the *in vivo* role of these gut-microbiota associated taxa is currently unclear. Here, we tackled this issue by examining the prevalence, origin, and activity of these taxa in the nasal microbiota of piglets.

**Results:** First, analysis of the nasal microbiota of farm piglets sampled in this study, as well as various publicly available data sets, revealed that gut-microbiota associated taxa indeed constitute a substantial fraction of the pig nasal microbiota that is highly variable across individual animals. Second, comparison of herd-matched nasal and rectal samples at amplicon sequencing variant (ASV) level showed that these taxa are largely shared in the nasal and rectal microbiota, suggesting a common origin driven presumably by the transfer of fecal matter. Third, surgical sampling of the inner nasal tract showed that gut-microbiota associated taxa are found throughout the nasal cavity, indicating that these taxa do not stem from contaminations introduced during sampling with conventional nasal swabs. Finally, analysis of cDNA from the 16S rRNA gene in these nasal samples indicated that gut-microbiota associated taxa are indeed active in the pig nasal cavity.

**Conclusion:** This study shows that gut-microbiota associated taxa are not only present, but also active, in the nasal cavity of domestic pigs, and paves the way for future efforts to elucidate the *in vivo* function of these taxa within the nasal microbiota.

## Introduction

The network of microorganisms inhabiting the bodies of animals is known as the microbiota [1–3]. The microbiota has been shown to play a pivotal role in various aspects of host health, for example by providing critical support in immune system maturation, nutrient utilization [2, 3], and defense against pathogen invasion [3, 4]. In the case of respiratory pathogens, one of the first lines of defense is the nasal microbiota [5], and many recent studies have focused on characterizing the commensal nasal microbiota and its relationship with respiratory pathogens in humans [5, 6] and various animal species [7–11]. These studies have identified a variety of taxa that are frequently found in the nasal microbiome of different host species, including members from different genera such as *Moraxella*, *Lactobacillus*, *Streptococcus*, *Haemophilus*/*Glaesserella*, and *Staphylococcus* [6–11].

Surprisingly, these studies also frequently detected microorganisms in the nasal cavity that are normally associated with the gut microbiota, in particular many anaerobic bacteria from the *Bacteroidales* and *Clostridiales* orders. These gut-microbiota associated taxa can be found in human nasal microbiota, where anaerobic Gram-negative bacteria such as *Prevotella* and *Veillonella* are frequently detected [6], but are particularly prevalent in the nasal microbiota of pigs [7]. Considering that the respiratory tract is unlikely to have anaerobic niches [12], there is ongoing discussion about the *in vivo* role of these gut-microbiota associated taxa in the nasal microbiota. Recent *in vivo* studies in piglets have shown that these taxa are differently abundant under different health-status scenarios [11, 13–17] and are variable through age stages [12, 18, 19], pointing towards a functional role. In contrast, other studies have attributed the presence of these gut-microbiota associated taxa to contamination from fecal material and/or soil [12, 20], which could be explained by the rooting behavior of pigs [12, 21, 22].

In this study, we investigate the presence of these gut-microbiota associated anaerobic taxa in the nasal microbiota of piglets. Specifically, we focus on three questions: **1)** how prevalent are these gut-microbiota associated taxa in the nasal microbiota of domestic pigs? **2)** what is their source (i.e. do these microorganisms truly originate from the gut)? and **3)** are these taxa active in the aerobic nasal environment? We tackle these questions using a combination of 16SrRNA amplicon sequencing of DNA (total communities) and cDNA retrotranscribed from RNA (active communities) in matched *in vivo* samples obtained from individual animals. We confirm that gut-microbiota associated taxa indeed represent a substantial fraction of the pig nasal microbiota across a wide range of samples from this study and literature. Comparison of Amplicon Sequencing Variants (ASVs) in matched rectal/nasal samples suggests a shared pool of these taxa in both body sites, pointing to a common source. Moreover, surgical sampling of the inner nasal tract indicates that these gut-microbiota associated taxa are not introduced during sampling but are truly located in the pig nose. Finally, comparison of total and active microbial communities suggests that these taxa are active throughout the nasal cavity of pigs. Overall, this work sheds light on the role of gut-microbiota associated taxa in the nasal microbiota of pigs and supports the notion that these taxa are not only present, but also active.

## Results

### Characterizing the gut-microbiota associated fraction in the pig nasal microbiota

As the starting point of this study, we aimed to characterize the fraction of gut-microbiota associated taxa found in the nasal microbial communities of piglets. Towards this end, we sampled the nasal cavity and rectum of 24 healthy animals from three different commercial farms located in Spain without a history of respiratory disease outbreaks and used 16S rRNA gene sequencing to determine the microbiota composition (**Figure 1A**). We obtained a total number of 9,103 different ASVs (mean of 132,051.44 read counts per sample) after processing the raw reads. To determine the fraction of gut-microbiota associated microorganisms in these samples, we focused on two orders (*Bacteroidales*/*Clostridiales*) which constitute the most abundant taxa in the gut microbiota and defined these as “gut-microbiota associated taxa”. We found that *Bacteroidales* and *Clostridiales* represented a substantial fraction of the pig nasal microbiota in most animals, accounting for 9.43 ± 3.8 % and 20 ± 7.7 % of the total composition, respectively (mean ± SD across samples) (**Figure 1B, left**). As expected, this fraction was higher in the respective rectal swabs (**Figure 1B, right**). We also determined this gut-microbiota associated fraction in the pig nasal microbiome in an alternative way by identifying taxa that are prevalent in a reference data set of gut microbiota (obtained from about 300 animals [23]) and obtained very similar results (**supplementary Text 1 and supplementary Figure 1**). Consistent with previous reports [7, 11, 12], in the nasal samples, *Prevotella* and *Bacteroides* together with *Ruminococcaceae* and *Lachnospiraceae,* were the most prevalent taxa within *Bacteroidales* and *Clostridiales*, respectively. Reassuringly, other reported nasal colonizers such as *Moraxella*, *Acinetobacter*, and *Enhydrobacter* genera from *Pseudomonadales*, *Lactobacillus* and *Streptococcus* genera from *Lactobacillales* and *Glaesserella* from *Pasteurellales* were also highly abundant in the nasal samples (**supplementary Figure 2**).

**Figure 1.**
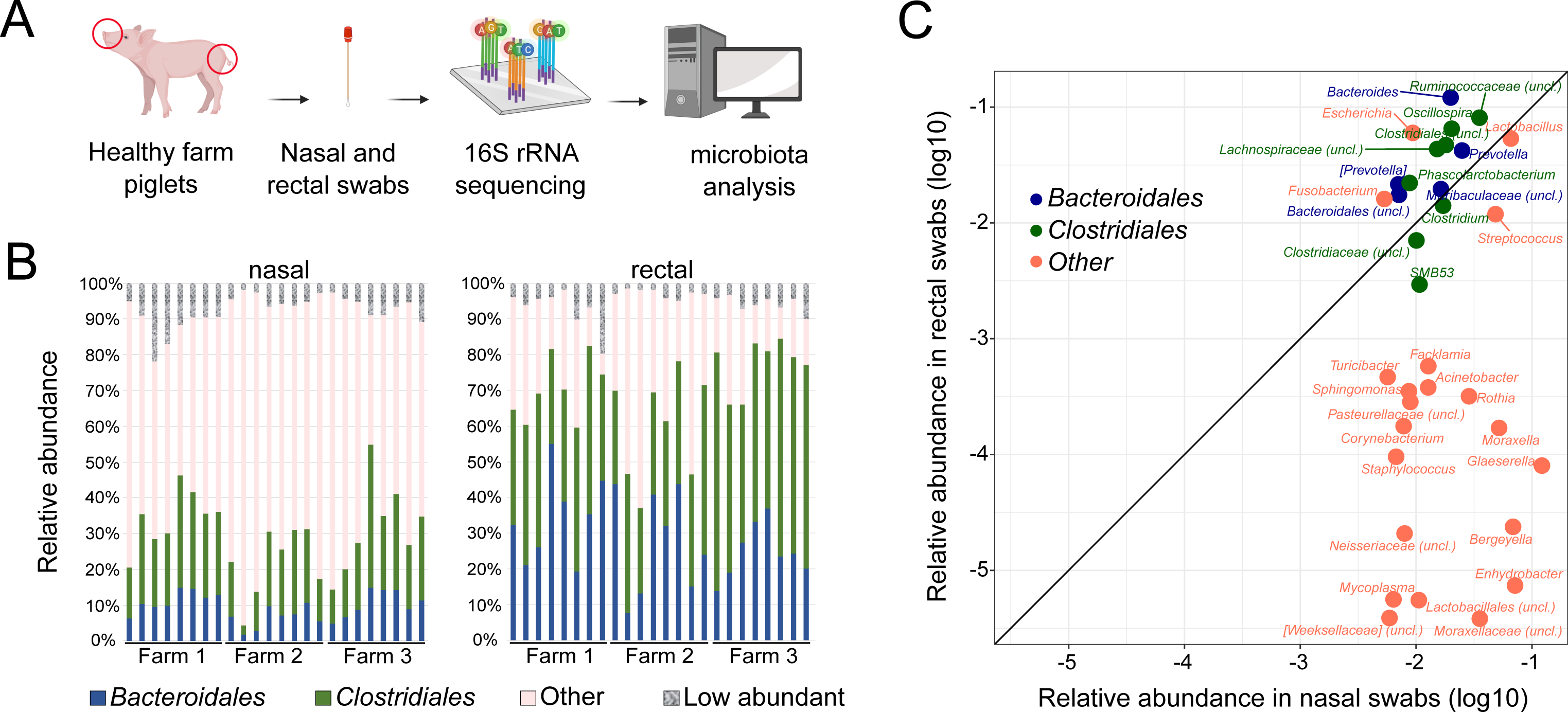
Detection of gut-microbiota associated taxa in nasal and rectal swabs from healthy 2–3 week-old piglets from three farms. **(A)** Schematic of microbiota sampling and sequencing approaches (created with BioRender.com). **(B)** Summed relative abundance of *Bacteroidales* (blue) and *Clostridiales* (green) taxa in nasal (left) and rectal (right) swabs of 24 individual animals. Summed abundance of other taxa (order level) with > 1% mean relative abundance across samples are shown as “Other” (pink). Orders with <1% relative abundance are summed in the category “Low abundant”. **(C)** Relative abundances in nasal and rectal samples (24 individuals mean) of genera in nasal microbiota with >0.5% mean relative abundance across animals. Square brackets in taxonomical assignations indicate contested names in the reference Greengenes database used (13.8).

To assess whether this substantial fraction of gut-microbiota associated taxa is a unique feature of these particular samples, we quantified the fraction of *Bacteroidales* and *Clostridiales* in 11 publicly available data sets of pig nasal microbiota samples ([11, 15, 20, 24–30], see **supplementary Table 1**). We found that across different countries of origin, pig ages, and sequenced regions, *Bacteroidales* and *Clostridiales* represent a substantial (albeit variable between individual animals) fraction of the nasal microbiota (**supplementary Figure 3, supplementary Table 2**). Taken together, these results confirm that the pig nasal microbiota has a substantial fraction of commonly gut-microbiota associated taxa.

### Identifying the source of gut-microbiota associated taxa in the pig nasal microbiota

The most probable source of gut-microbiota associated taxa in the pig is the gut microbiota itself, and in support of this hypothesis we found that the most abundant genera belonging to either *Bacteroidales* or *Clostridiales* in rectal samples also tended to be highly abundant in matched nasal samples (**Figure 1C**). On the other hand, most genera previously reported as nasal colonizers exhibited low abundances in rectal samples (**Figure 1C)**. However, it is conceivable that although the gut-microbiota associated taxa in these two body sites belong to the same genera, pig nose and gut may nevertheless be inhabited by distinct strains with different niche preferences.

To address this question, we examined the individual ASVs (as a proxy for strain identity [31]) found in nasal and rectal swabs within each farm. Specifically, for each farm we identified the most abundant *Bacteroidales* and *Clostridiales* ASVs in nasal and rectal samples and determined their overlap (the 100 most relatively abundant ASVs in nasal and rectal samples were selected, **Figure 2A, supplementary Table 3**). We found that 39%, 14% and 30% of ASVs belonging to *Bacteroidales,* were shared between nose and rectal samples in farms 1, 2 and 3, respectively. In the case of *Clostridiales,* these proportions were 36%, 9% and 39%. Importantly, we found that the ASVs shared between both body sites were often more highly abundant than the site-specific ones (especially in farms 1 and 3). Moreover, we did not observe any biases for certain bacterial families or genera within the body-site specific ASVs. For example, we found ASVs classified as *Prevotella*, *Bacteroides*, *Veillonella* and *Oscillospira* among the site-specific ASVs in both nasal and rectal samples (**supplementary Table 3**). In contrast, ASVs belonging to families of known nasal colonizers such as *Moraxellaceae*, *Pasteurellaceae* and *Weeksellaceae* showed a much lower degree of overlap between body sites (**Figure 2B**), as most of their ASVs were present in nasal but largely absent in rectal samples. Among the exceptions we found *Lactobacillaceae* and *Streptococcaceae*, whose ASVs still exhibited substantial overlap between the two body sites. Thus, the analysis of nasal and rectal microbiota at ASV level suggests a shared origin of the gut-microbiota associated taxa (presumably environmental e.g. through transfer of fecal matter) found in the pig nasal and rectal microbiota.

**Figure 2.**
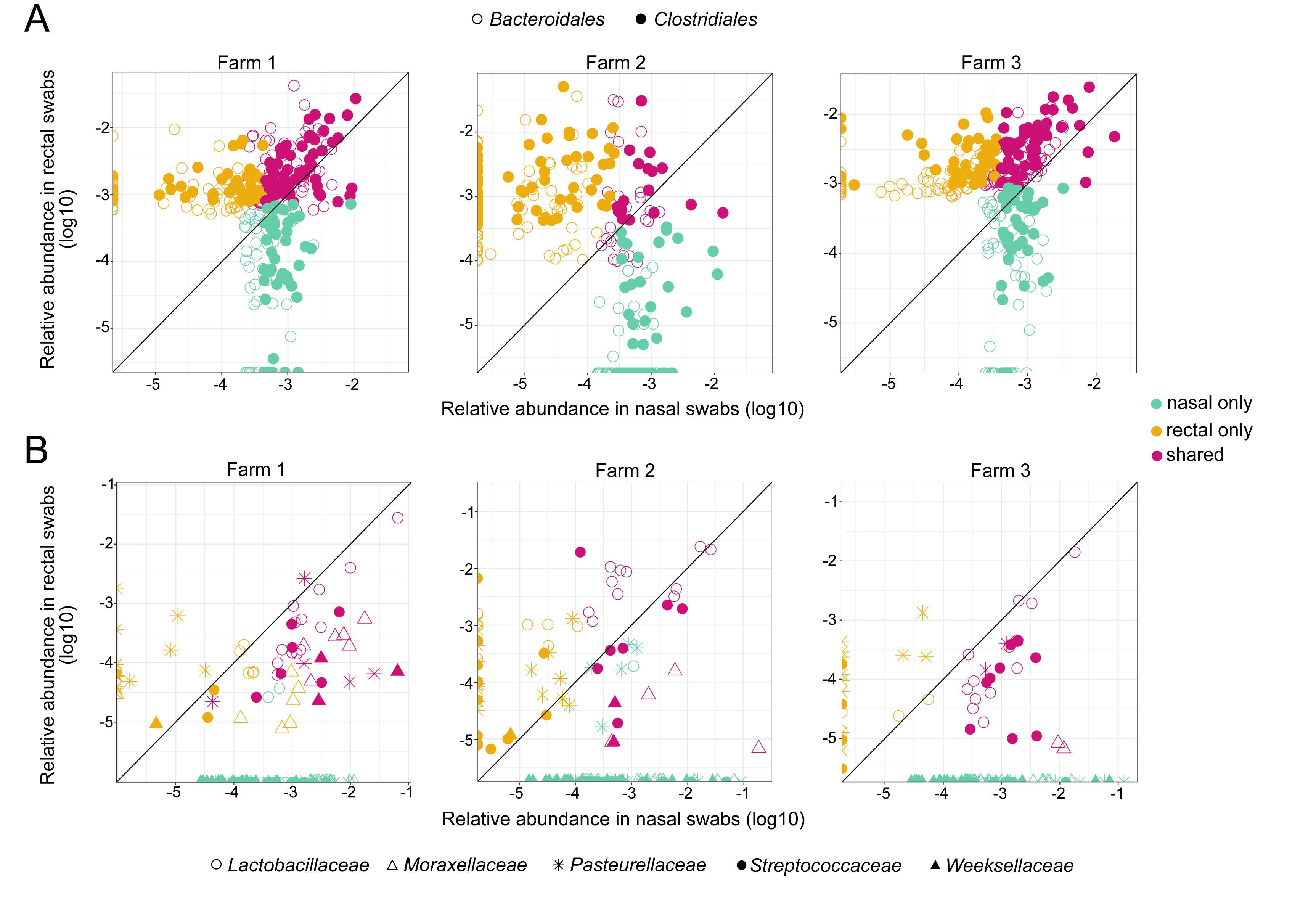
Nasal and rectal overlap of gut-microbiota associated taxa (A) and nasal colonizers (B) most prevalent ASVs per sampled farm. For gut-microbiota associated taxa the 100 most abundant ASVs in nasal and rectal samples were selected. For other nasal colonizers’ families (**B**), the top 20 ASVs were considered. Each dot corresponds to one ASV (mean abundance of 8 animals from each herd). Turquoise: ASV is among the most abundant ASVs in nasal but not rectal swabs. Yellow: ASV is among the most abundant ASVs in rectal but not nasal swabs. Pink: ASV is among the most abundant ASVs in both sites.

### Surgical deep sampling of the pig nasal cavity to identify gut-microbiome associated taxa at different depths

The shared gut-microbiota associated taxa in nasal and rectal swabs described above are consistent with the hypothesis that these taxa enter the pig nasal cavity through the transfer of fecal matter driven by the animals’ rooting behavior. To further validate that these taxa were indeed resident in the pig nasal cavity and did not stem from contaminations introduced by the sampling through the nostrils, we performed additional in-depth sampling (**Figure 3A**). Specifically, we surgically opened the nasal cavity of five animals post-mortem to enable sampling of its (normally inaccessible) deep and middle parts (as detailed in the Methods section), avoiding a possible swab contamination from the skin surrounding the nostril openings. Additionally, standard nasal swabs, swabs of the external nasal area, and rectal swabs of each animal were taken for comparison. We observed a gradient in microbial load (highest to lowest, as determined by 16S rRNA gene qPCR) from external to deep internal samples, which was consistent with the number of total reads and abundance of contaminant sequences from negative control samples (**supplementary Figure 4**). Moreover, the deep nasal microbiota exhibited lower richness compared to samples from external nose (Chao1 index *p* < 0.05, **supplementary Figure 5A**), and was identified as a distinct community (as determined by beta diversity analysis) compared to the external nares (Jaccard and Bray-Curtis PERMANOVA *p* < 0.05, **supplementary Figure 5A, supplementary Table 4**), when deep nasal samples were compared pairwise with standard and external swab samples. Comparison of the composition at different nasal cavity locations within the same animal revealed that those taxa which were found at all sampling sites (of which there were only few in any given animal) tended to constitute the bulk of the observed microbiota (**supplementary Figure 6, see supplementary text 2 for detailed compositional analysis**). This dominance of few taxa was found both at the genus level, as well as at the level of individual ASVs, suggesting that the nasal microbiota of individual animals is consistently composed of few dominating strains across the whole nasal cavity. Importantly, nasal swabs as well as matched surgical nasal samples showed a variable, but substantial, fraction of gut-microbiota associated taxa (**Figure 3B and supplementary Figure 5B**), many of which were found consistently throughout the nasal cavity in each animal (**Figure 3C**). Thus, these findings suggest that gut-microbiota associated taxa do reside in the pig nasal cavity and do not stem from contaminations introduced by the sampling procedure itself.

**Figure 3.**
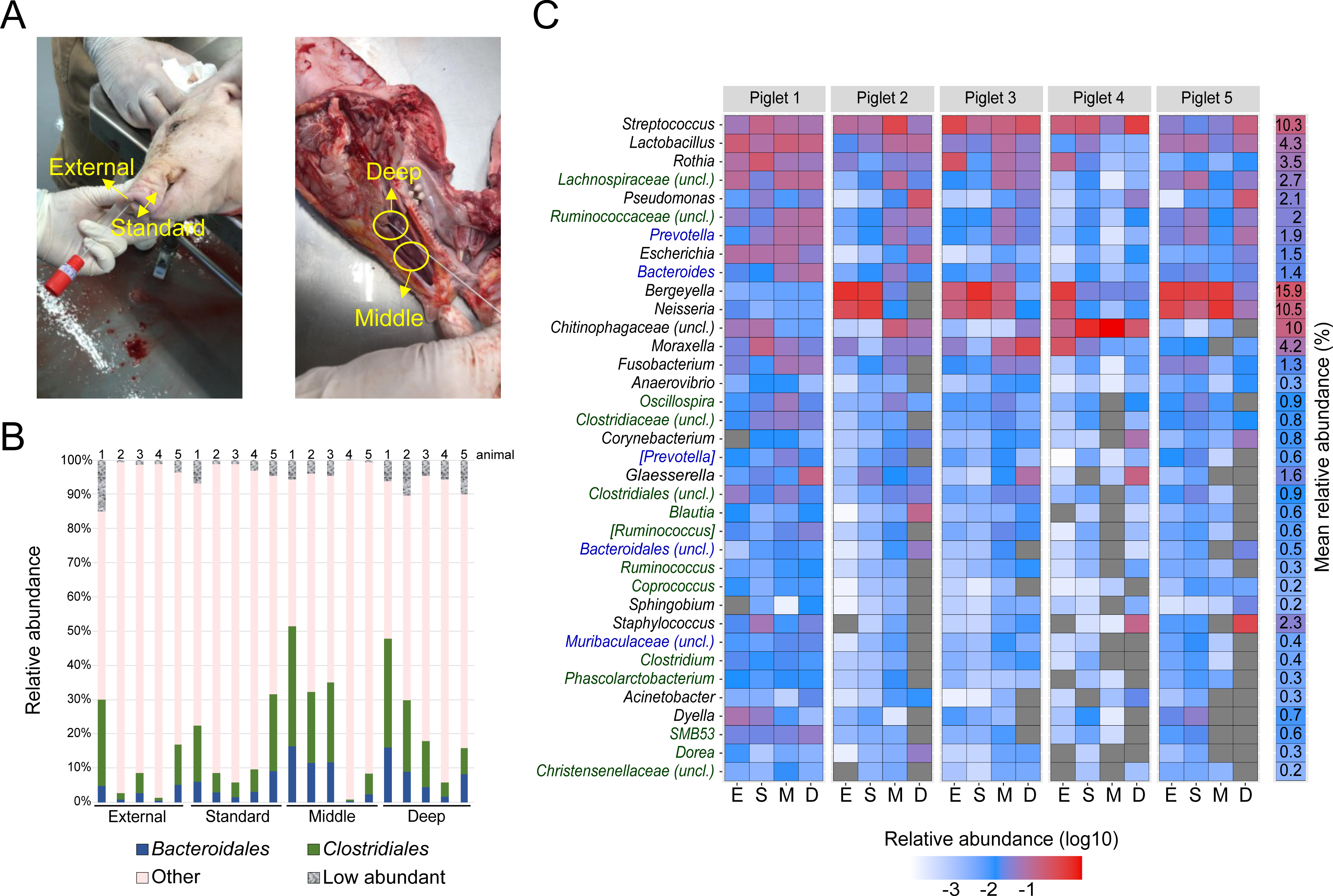
Characterization of surgical microbiota samples. **(A)** Collection of nasal samples at four sites. External and standard nasal sampling; deep and middle nose samples after longitudinal surgical cuts of piglet heads (see methods). **(B)** Summed relative abundance of *Bacteroidales* (blue) and *Clostridiales* (green) taxa in the different types of nasal swabs of the 5 individual animals. Summed abundance of orders with > 1% mean relative abundance across samples are shown as “Other” (pink). Taxa with <1% relative abundance are summed in the category “Low abundant”. **(C)** Most prevalent genera across the nose of the 5 sampled piglets. Genera are ordered from top to bottom by prevalence (present in most samples) and global relative abundance. Gut-microbiota associated genera are labelled in blue (*Bacteroidales*) and green (*Clostridiales*). Numbered column on the right: mean relative abundance across all samples. Sample sites are marked as E(xternal), S(tandard), M(iddle), D(eep).

### Assessing the activity of gut-microbiota associated taxa by quantifying the 16S rRNA transcripts

The results described above suggested that gut-microbiota associated taxa are present throughout the whole pig nasal cavity. Next, we wanted to determine whether these taxa (which include many obligate anaerobic species) are also active in the aerobic environment of the pig nose. Towards this end, we quantified 16S rRNA gene transcripts (a proxy for protein synthesis potential and thus indirect measure of cellular activity, [32–34]) in the nasal microbiota samples described above. We found that gut-microbiota associated taxa constituted a similar portion in these RNA derived samples as in the respective DNA samples described above. Specifically, in nasal samples taken from 24 animals across 3 farms, *Bacteroidales* and *Clostridiales* orders accounted for a mean 7.1% ± 6.2 and 22.4% ± 14.1, respectively (**Figure 4A and Supplementary Figure 7A**). In the surgical samples taken at different sites in the nasal cavity, *Bacteroidales* accounted for 10.3% ± 8.8 (Deep), 8.2% ± 10.4 (Middle), 1.6% ± 3.2 (Standard) and 0.1% ± 0.1 (External); and *Clostridiales* for 3.6% ± 2.8 (Deep), 3.9% ± 3.6 (Middle), 3.2% ± 5.6 (Standard) and 2.9% ± 4.6 (External) (**Figure 4A and Supplementary Figure 7B**). Moreover, we found that gut-microbiota associated taxa had similar RNA to DNA ratios as the ratios for reported nasal colonizers when examined globally (summing across different taxa) in individual animals (**Figure 4B**), or when examining different genera (**Figure 4C**) and families (**supplementary Figure 8**). To identify individual families belonging to gut-microbiota associated taxa that deviate from this general trend, we finally compared their RNA/DNA ratio distributions with those of nasal colonizers in the farm animal samples (**supplementary Figure 9**). With some exceptions (e.g. lower RNA/DNA ratios for *Bacteroidales (unclass.)* and higher ratios for *Clostridiaceae*), RNA/DNA ratio distributions were largely not significantly different for most gut-microbiota associated taxa compared to reported nasal colonizers. Taken together, these results suggest that gut-microbiota associated taxa are not only present in the pig nasal environment, but also active.

**Figure 4.**
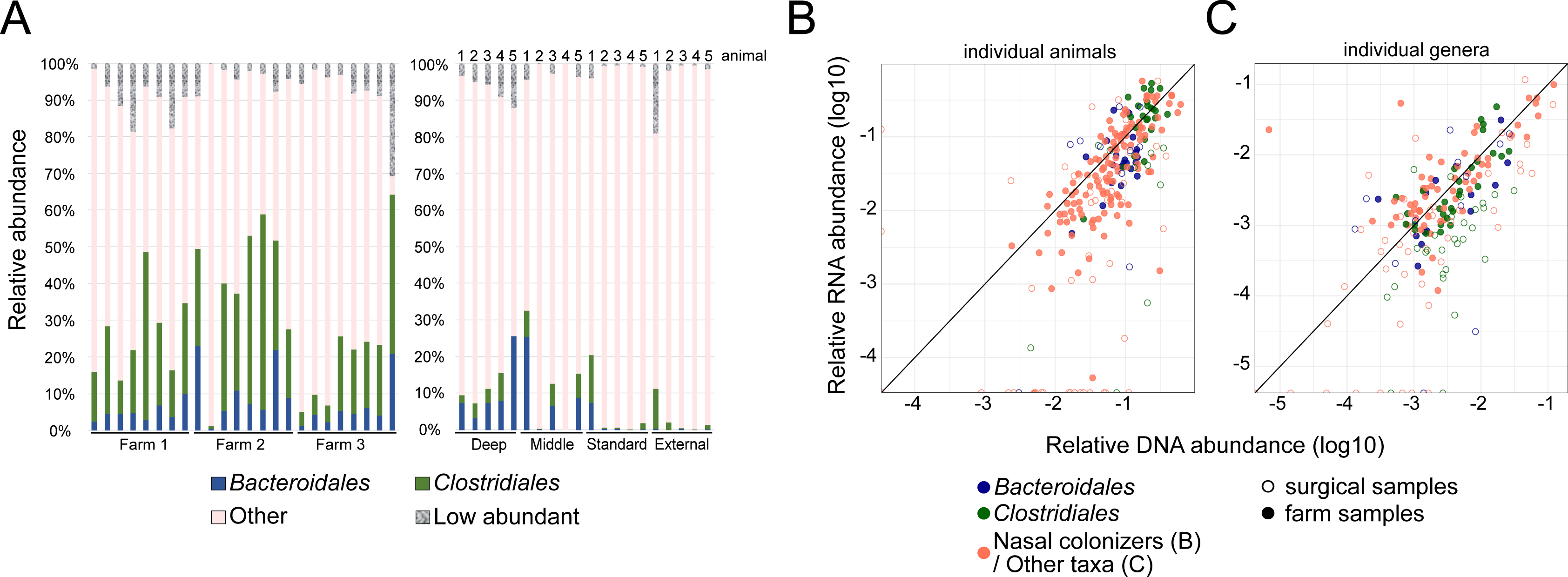
RNA-based quantification of nasal microbiota activity. **(A)** Summed relative abundance of *Bacteroidales* (blue) and *Clostridiales* (green) taxa in the two sets of nasal samples (nasal swabs from 24 farm animals, left; surgical samples from 5 animals, right; see methods). Other taxa (order level) with > 1% mean relative abundance are shown as “Other” (pink). Taxa with <1% relative abundance are summed in the category “Low abundant”). **(B)** Relative abundance in RNA/DNA samples of Bacteroidales (blue), Clostridiales (green), and nasal colonizers (*Lactobacillaceae*, *Moraxellaceae*, *Pasteurellaceae*, *Streptococcaceae* and *[Weeksellaceae]*, pink). Each dot corresponds to the abundances of the mentioned taxa in one individual animal. Open circles: invasive nasal samples from 5 animals (deep and middle nasal cavity, see Figure 3). Filled circles: standard nasal swabs from 24 animals across 3 farms. Shown are mean values across all samples. **(C)** Relative abundance in DNA and RNA samples (as determined by 16S rRNA sequencing of DNA and cDNA, see main text) of the most relatively abundant genera, selected as those > 0.1% mean abundance in DNA or RNA farm samples. Gut-microbiota associated taxa (*Bacteroidales* and *Clostridiales*) are labelled in blue and green, respectively. Genera from other orders are shown in pink. Open circles: deep nasal samples obtained surgically from 5 animals (deep and middle nasal cavity, see Figure 3). Filled circles: standard nasal swabs from 24 animals across 3 farms. Shown are mean values across all samples

## Discussion

In this study, we aimed to elucidate the role of typically gut-microbiota associated taxa in the nasal cavity of domestic pigs. Specifically, we asked three questions: 1) how prevalent are these taxa *in vivo* and across different anatomical sites in the nasal cavity?; 2) do these taxa indeed stem from the gut microbiota?; and 3) are these taxa active in the pig nasal cavity? To answer these questions, we used a combination of regular swab and surgical deep sampling of the pig nasal microbiota and inferred its composition/activity by 16S rRNA gene DNA and cDNA sequencing, respectively. These efforts yielded three key findings.

First, we found that gut-microbiota associated taxa constitute a substantial fraction of the nasal microbiota both across individual animals, as well as across different sites within the nasal cavity. This finding is in agreement with previous studies, which sampled the pig nasal microbiota, and is also consistent with other studies relying on sampling of the lower respiratory tract of pigs, where *Clostridium* and *Prevotella* genera were prevalent [7, 13, 35]. Thus, our results confirm that gut-microbiota associated taxa are part of the swine nasal respiratory microbiota and are not a sampling artifact. Nevertheless, the sampling of environments that do not have high microbial biomass, including many anatomical sites in the respiratory tract [36, 37], can pose technical challenges such as the false detection of transient environmental microbes [37]. Although we controlled for some potential sources of contamination (sequencing blank controls and getting undisturbed nasal samples) more studies are needed to validate the inhabitants of the pig nasal microbiota.

Second, our analysis of matched nasal and rectal microbiota samples from different farms showed a large overlap between gut-microbiota associated taxa from these two sites at ASV level. This finding suggests a common source of these taxa in both anatomic sites, for example in the form of solid fecal matter that enters the nasal cavity [12, 20], or through the air within a farm. In support of this hypothesis, a recent study found that gut-microbiota associated taxa are highly abundant in the air of pig farms, as well as in the nasal microbiota of pig farmers (compared to non-exposed individuals) [38, 39], suggesting substantial flow of bacterial material within animal farms. In contrast to gut-microbiota associated taxa, many other dominant taxa in the nasal microbiota, such as *Moraxellaceae*, *Pasteurellaceae,* and *Weeksellaceae*, showed a low degree of overlap between the two body sites, indicating that they may be professional nasal colonizers with reduced ability to colonize other (i.e. anaerobic) niches. Interesting exceptions were the *Lactobacillus* and *Streptococcus* genera, which also overlapped substantially at ASV level between these two body sites. Given that both *Lactobacillus* and *Streptococcus* are facultative anaerobes, it is tempting to speculate whether these genera may in fact predominantly reside in the gut, but are also able to colonize the nasal cavity (and possibly other sites) if presented with the opportunity.

Third, our quantification of 16S rRNA expression showed that gut-microbiota associated taxa are indeed active in the pig nasal cavity. To our knowledge, this is the first study which quantified pig nasal microbiota activity *in vivo*, and it suggests that these gut-microbiota associated taxa are not merely inactive transient members in the nasal cavity. Notably, recent metatranscriptomics analyses did report expression of some gut-microbiota associated taxa (e.g. *Prevotella*) in nasal samples taken from children [40] and adult patients with asthma [41], suggesting that these taxa may also be active in the human nasal cavity. One caveat is that we quantified 16S rRNA gene transcripts to determine the activity of gut-microbiota associated taxa in the pig nasal microbiota, which is an imperfect measure of cellular activity [32]. Future efforts could examine the activity of these taxa with methods that quantify metabolic activity more directly, for example by using fluorescently-tagged metabolic probes [42–44].

This study has several limitations. First, we sampled animals at an early age (i.e. pre-weaning stage at 2-3 weeks of age). Our choice was motivated by the fact that pigs at this age are most susceptible to respiratory pathogens, while the nasal microbiota is still rather unstable [11, 45]. Nevertheless, our analysis of other published studies did reveal that gut-microbiota associated taxa are also prevalent at later stages in the animals’ life [20, 25, 29], and future studies may examine to which extent the findings presented herein also hold in adult pigs and other species.

Second, although our surgical samples suggest that gut-microbiota associated taxa are present throughout the pig nasal cavity, the methods used in this study are not able to resolve whether these taxa are located in any specific anatomical niches. Scanning electron microscopy imaging of the upper respiratory tract of piglets has recently suggested the absence of anaerobic crypts [12], but it is conceivable that there may still be anaerobic substructures e.g. within or under the mucosal layer. Such potential substructures or locations may explain why strictly anaerobic bacterial taxa survive and are active in a basically aerobic environment such as nasal cavity. To resolve these questions, future efforts may use *in vivo* imaging methods like these [46, 47] to resolve the microstructure of the nasal microbiota in more detail.

Finally, while our results do suggest that gut-microbiota associated taxa are indeed active in the pig nasal cavity, we did not examine the *in vivo* function of these taxa within the pig nasal microbiota. Recent *in vivo* studies [11, 13–17, 48] have shown that the presence of these taxa is associated with disease outcomes and dysbiosis in various farm animals, but also in humans [41]. For example, in humans with chronic rhinosinusitis, *Prevotella* (genus belonging to *Bacteroidales*) was associated with increased inflammatory severity [49]. Future studies may use the results presented here as a starting point to elucidate the mechanistic underpinnings of these associations.

In conclusion, this study suggests that gut-microbiota associated taxa are indeed present and active in the nasal cavity of domestic pigs. These findings may serve as a starting point for future research aiming at elucidating the *in vivo* function of these taxa within the nasal microbiota.

## Methods

### Animal experimentation and ethics approval

Animal experimentation was performed following proper veterinary practices, in accordance with European (Directive 2010/63/EU) and Spanish (Real Decreto 53/2013) regulation. Sampling in farms was done with the approval of the Ethics Commission in Animal Experimentation of the Generalitat de Catalunya (Protocol number 11213). For the surgical sampling of five animals’ different nasal depths, euthanasia was performed following good veterinary practices. According to European (Directive 2010/63/EU of the European Parliament and of the Council of 22 September 2010 on the protection of animals used for scientific purposes) and Spanish (*Real Decreto* 53/2013) normative, this latter procedure did not require specific approval by an Ethical Committee (Chapter I, Article 3. 1 of 2010/63/EU).

### Sample collection

*Farm Samples*: Matched nasal and rectal swabs were taken from 24 healthy 2-3-week-old piglets from three Spanish farms without reported respiratory diseases (termed Farm 1, Farm 2, Farm 3). After sampling, swabs were placed into sterile tubes filled with 1000µL DNA/RNA shield (Zymo Research) and transported to the laboratory under refrigeration.

*Surgical Samples*: Deep surgical samples of the nasal cavity were obtained as follows: five healthy piglets of 2-3 weeks of life were moved to IRTA-CReSA facilities and euthanized by means of an overdose of sodium pentobarbital (Dolethal®). Four types of nasal swabs per piglet were taken (labeled standard, external, middle, and deep) as described below. First, a nasal swab from one nostril was taken (“standard” swab, Figure 3A). Second, an external swab was taken by introducing the swab superficially in the second nostril. Afterwards, deep and middle swabs were taken (at positions outlined in Figure 3A) from the second nostril after longitudinally cutting and separating the skin of each piglet’s head (to prevent contaminations from the skin surface) and subsequent cutting the skull, directly from the nasal turbinate without touching any other part of the piglet nose to avoid contamination and after removing the cartilaginous nasal wall. Additionally, rectal swabs were obtained from the same animals. Sample swabs were collected in sterile plastic tubes filled with 800µL of DNA/RNA shield (Zymo Research) to ensure the stability and preservation of the genetic material. Negative controls (sterile DNA/RNA shield without a swap) were included in each extraction. Samples were stored at -20°C until extraction.

### DNA/RNA extraction

Metagenomic DNA and RNA were extracted starting from 350 µL of DNA/RNA shield swab sample (previously vortexed) and following a modified protocol of ZymoBIOMICS DNA/RNA Miniprep kit (Zymo Research) in which the lysis was performed only chemically using 700 µL of ZymoBIOMICS DNA/RNA lysis buffer (2 volumes of lysis buffer per 1 volume of sample). The RNA fraction of the samples was treated with 80 µL of DNAse I (included in the kit) at room temperature for 20 minutes. Elution of both DNA and RNA was done in 50 µL of elution buffer. DNA and RNA concentration was measured using BioDrop DUO (BioDrop Ltd). DNA and RNA were stored at -80°C until sequencing.

### 16S rRNA Sequencing

16S rRNA gene libraries were prepared from the total extracted DNA and cDNA from RNA samples and sequenced at *Servei de Genòmica, Universitat Autònoma de Barcelona* (Illumina pair-end 2X300 bp, MS-102-2003 MiSeq Re-agent Kit v2, 500 cycle), using Illumina recommended primers for variable regions V3V4 of 16S rRNA gene (fwd 5’TCGTCGGCAGCGTCAGATGTGTATAAGAGACAGCCTACGGGNGGCWGCAG, rev 5’ GTCTCGTGGGCTCGGAGATGTGTATAAGAGACAGGACTACHVGGGTATCTAATCC). The size of the amplicons was verified on a Bioanalyzer DNA 1000 chip (Agilent), as expected amplicon length was approximately 460bp. Finally, the sequences were sorted into samples and used as input for bioinformatic analysis.

### Bioinformatic analysis of 16S rRNA sequencing data

The microbiota composition of the samples was analyzed with quantitative insights into microbial ecology (QIIME) 2 software version 2022.2 [50]. The detailed pipeline followed from raw reads to obtain tables of abundances at ASV and other taxonomic levels, the diversity analyses and the statistical tests to determine the differential abundances, can be found at https://zenodo.org/record/8013997 (Zenodo ID 8013997). Briefly, the pre-processing of the reads was done separately for each sequencing run. At first, raw demultiplexed reads with quality (fastq) were imported and their quality was evaluated with *qiime2 demux* plugin. Primers from the variable region V3-V4 were extracted using *qiime2 cutadapt* plugin [51] Sequences that did not contain primers and thus, were not a sequencing product, were removed from the analysis. DADA2 software package [52] was used under the parameters detailed in the pipeline to quality filter, paired-end merge, remove chimeras and sort reads into ASVs. Contaminant artifactual amplicons from non-prokaryotic origin were identified with *qiime2 quality control* plugin [53] by aligning with VSEARCH [54] all ASVs against Greengenes database Vs. 13.8 [55] clustered with 88% identity (available at https://docs.qiime2.org/2022.2/data-resources/). Unmatched sequences were filtered out from the analysis. Additionally, ASVs identified as *Archea*, *Mitochondria* or *Chloroplast* (which also contain 16S rRNA), were also discarded. The alignment of the remaining sequences was performed with MAFFT [56], and the hypervariable positions were masked [57] with *qiime2 alignment* plugin. Finally, ASVs found in the negative control of the deep nose dataset were removed from the analysis (21 and 41 for DNA and RNA controls, respectively). The taxonomic classification of the ASVs was performed using a naïve Bayes classifier with scikit-learn Python module for machine learning [58]. In order to increase the classifier accuracy [59], it was previously trained against prokaryotic 16S rRNA gene V3-V4 region, extracted from Greengenes database (13.8 version) clustered at 99% sequence identity. This qiime2 feature classifier artifact can be found at https://zenodo.org/record/8013997 (Zenodo ID 8013997). Square brackets in taxonomical assignations indicate contested names in the reference Greengenes database used (13.8).

In order to normalize uneven sampling depths [60], the diversity analyses of the samples from the surgical sampling (supplementary text 2) were performed at a normalized depth of 2291, corresponding to the lowest depth sample. The alpha diversity of the samples was estimated with Chao1 [61] and Shannon [62] indexes. Significant differences were found with pairwise non-parametric t-tests (999 random permutations) using *qiime2 diversity alpha-group-significance* plugin [63]. Beta diversity was calculated with Jaccard [64] and Bray-Curtis [65] dissimilarity indexes for the qualitative and quantitative analyses, respectively. *Qiime2 core-metrics* plugin [66, 67] was used to compute principal coordinate (PCoA) analysis. The percentage of explanation of the variables under study was estimated with the Adonis function from the Vegan package, in R software [68]. The significance of beta diversity analyses was calculated by PERMANOVA pairwise test (999 random permutations) using *qiime2 diversity beta-group-significance* plugin [69]. Differently abundant taxa between groups were found with ANCOM-BC algorithm [70]. For all the stated analyses, the significance threshold *p* value was set to 0.05. Output microbiome data processing and plot generation was performed using R script language version 4.2.2 in RStudio environment version 2022.07.0[71], using the packages qiime2r [72], reshape2 [73], ggplot2 [74] and tidyverse [75], as well as MATLAB (version 2021A).

### Quantification of total 16S rRNA gene concentration

Total 16S rRNA gene concentrations (as a proxy for microbial load in swabs) were quantified as follows. Briefly, the reaction was prepared in a volume of 20 µL consisting in 2 µL of the template DNA and 18 µL of Femto Bacterial qPCR Premix (Femto Bacterial DNA Quantification Kit, Zymo Research). The PCR reaction was performed using a 7500 Fast Real-Time PCR System (ThermoFisher Scientific) at 95°C for 10 min, 40 cycles of 95°C for 30s, 50°C for 30s, 72°C for 1 min, followed by a melting curve and a final extension of 72°C for 7 min. Each run contained two replicates per sample, a standard curve of 6 points (2 to 0.00002 ng), a negative and a positive extraction control as well as PCR negative controls. Graphpad 8.3 (538) Prism software (Dotmatics, San Diego CA) was used to analyze the data obtained from the 16s rRNA qPCR of DNA extracted from the swabs.

## Data availability

The raw data used in this study are publicly available at NCBI’s SRA database under BioProject ID PRJNA981084. Processed data sets are available as supplementary tables 1-5 (see descriptions below).

## Supplementary information

Supplementary text and figures are provided as a single document containing:

- **Supplementary text 1** – identification of gut-microbiota associated taxa in the pig nasal microbiota
- **Supplementary text 2** – compositional analysis of deep pig nasal microbiota samples
- **Supplementary Figures 1 – 9**

Supplementary tables are provided as individual files as described below:

- **Supplementary Table 1** – List of publicly available data sets used in this study.
- **Supplementary Table 2** – Relative abundance of gut-microbiota associated taxa (i.e. *Clostridiales* and *Bacteroidales* orders) across individual samples in 11 publicly available data sets used in this study.
- **Supplementary Table 3** – Most relatively abundant gut-microbiota associated ASVs in nasal and rectal samples (top 100 in each body site) in each of the three sampled farms.
- **Supplementary Table 4** – ANCOM-BC statistically significant (corrected q-value < 0.05) differences between sampling sites in nasal cavity at ASV, order, family, genus and species taxonomic levels.
- **Supplementary Table 5** – Relative abundance of all identified taxa at ASV, order, family, genus and species level across the two groups of samples used in this study.
- **Supplementary Table 6** – Sample metadata file containing identifiers and description of the samples used in this study.

## Acknowledgements

The authors are grateful to Nuria Galofré-Milà and Eva Huerta for their technical support. The authors are also grateful to the Centres de recerca de Catalunya (CERCA) Program.

## Funding

This work was supported by funding from the Spanish Ministry of Research and Innovation (RYC2021-033035-I to KK, PID2019-106233RB-I00/AEI/10.13039/501100011033 to VA and FCF). LBL and POG are supported by FPI(PRE2020-096048) and FPU (FPU19/02126) fellowships, respectively.

## Author contributions

Conceived and designed the study: KK, FCF, VA. Performed experiments: LBL, MS, JS. Performed analyses: POG, LBL, KK. Supervised the analyses: KK and FCF. Wrote the manuscript with contributions from all authors: POG, KK.

## Conflict of interest

The authors declare no conflict of interest.

## Notes

https://www.ncbi.nlm.nih.gov/bioproject/PRJNA981084

### Competing Interest Statement

The authors have declared no competing interest.

