## Supplementary Information for "Gut-associated microbes are present and active in the pig nasal cavity"

*: shared-first authors

#: shared-last authors

### Contents

Contents ... 1
Supplementary text ... 2
Supplementary text 1 – identifying gut-associated taxa in the pig nasal microbiome ... 2
Supplementary text 2 – compositional analysis of invasive pig nasal microbiome samples ... 3
Supplementary Figures ... 5
References ... 14

### Supplementary text

#### Supplementary text 1 – identifying gut-associated taxa in the pig nasal microbiome

To identify taxa in the pig nasal microbiome that may potentially stem from fecal contamination, we initially focused on two orders, namely *Bacteroidales* and *Clostridiales*. These two orders represent the largest fractions of the gut microbiome in humans and pigs. Moreover, at least *Clostridiales* are considered to be obligate anaerobes (and *Bacteroidales* include obligate and facultative anaerobes), and therefore are not expected to thrive in the largely aerobic environment of the nasal cavity (see e.g., [1] for $O_2$ levels in human respiratory tract).

To test whether the pig nasal microbiota contains other potentially gut-microbiome associated taxa not included in these two orders, we used a reference gut microbiome study from Xiao et al [2] and identified taxa that are frequently found in the microbiome of 288 healthy piglets sampled in different countries. We identified 36 taxonomical families that are present (relative abundance >= 0.01%) in at least 10% of the animals (using processed data obtained from http://gigadb.org/dataset/view/id/100187). We used these low thresholds to include also families with low prevalence. Of these taxonomical families, 33 were also detected in the pig nasal microbiota estimated from 24 farm samples. Four of these families (*Streptococcaceae, Staphylococcaceae, Lactobacillaceae, Pasteurellaceae*) represent well-established pig nasal commensals and were excluded from further analyses. We found that the summed relative abundance of these more refined gut-associated taxa in the nasal microbiome samples was very similar to the metric used above (i.e., labeling all *Clostridiales/Bacteroidales* as gut-microbiota associated), and was similarly dominated by *Clostridiales* and *Bacteroidales* (**Supplementary Figure 1**). Moreover, some taxa found in our samples belonging to these two orders could not be compared to these reference gut-microbiome taxa due to unresolved classification at the family level. To avoid potential biases by such unresolved classification at the family level, we decided to use *Clostridiales* and *Bacteroidales* orders as our indicators of potential gut-microbiota associated taxa throughout this study.

**Supplementary text 2 – compositional analysis of invasive pig nasal microbiome samples**

During the characterization of the different nose depth samples of the five animals necropsied, the bacterial load, measured by 16S rRNA gene concentration (ng/µL), was detected in a decreasing gradient from outside to inside the nasal cavity (Mann-Whitney test $p < 0.05$, **Supplementary Figure 4A**). The external nasal swabs contained the highest concentration of all nasal samples (mean of 0.081 ± 0.205 ng/µL), followed by the standard nasal swabs (0.015 ± 0.014 ng/µL), the middle nasal swabs (0.003 ± 0.002 ng/µL) and finally, the deep nasal swabs (0.001 ± 0.001 ng/µL). The concentration of bacteria was even higher in rectal samples (2.211 ± 1.986 ng/µL). Similarly, the number of reads obtained from 16S rRNA gene sequencing from each type of samples also decreased through the depth of the nose, while the representation of the ASVs found in the negative control correlated inversely as these represented a mean relative abundance of 18.3%, 3.2%, 1.5%, 0.6% and 0% in deep, middle, standard, external and rectal samples. (**Supplementary Figure 4B**).

When the diversity between the five types of samples was compared (beta diversity) a strong effect size was found in both qualitative (23%) and quantitative (29%) analyses (Adonis function $R^2$, $p < 0.05$), as rectal samples formed a differential cluster. In order to exclusively compare the different types of nasal samples, we excluded the rectal samples from the diversity analysis. Interestingly, despite some individual pairwise differences, the quantitative beta diversity analysis became non-significant statistically (PERMANOVA $p > 0.05$). On the contrary, the qualitative analysis still explained the 19% of group differences (Adonis test R2, $p = 0.003$). In the pairwise analysis, we detected that most differences occurred between the deep nose and the external samples in both qualitative and quantitative analyses (**Supplementary Figure 5A**). Regarding the alpha diversity, there were no differences between nose locations (Shannon index $p > 0.05$), but deep nasal samples reported a lower species richness than the external and standard nasal samples (Chao1 index, **Supplementary Figure 5A**, $p < 0.05$),

In order to characterize the most prevalent bacteria in the different parts of the swine nasal tract, we focused in the most dominant taxa (**Supplementary Figure 5B**). The most abundant taxa in the deep nose belonged to the orders *Lactobacillales*, such as *Streptococcus* and *Lactobacillus*; *Clostridiales* (composed of lower abundant genera within the families *Ruminococcaceae* and *Lachnospiraceae)*; and *Pseudomonadales*, mainly represented by *Moraxella* and *Pseudomonas*. Other relatively abundant genera were *Haemophilus*, *Chitinophagaceae* (*uncl*.) and *Staphylococcus*. Samples from the middle nose were dominated by *Clostridiales* (with a similar composition of many low abundant genera as the deep nose), *Lactobacillales* (mainly *Streptococcus* and *Lactobacillus*) and *Bacteroidales*, with *Prevotella* and *Bacteroides* as the most abundant. *Neisseria* was the dominating genus in only one sample. One sample was fully composed of the *Chitinophagaceae* (*uncl*.) genus (94%). The most relatively abundant taxa in the samples from the external nose were *Neisseria*, *Streptococcus*, *Rothia*, *Enhydrobacter* (*Pseudomonadales*), *Moraxella* and *Lactobacillus*, with their respective orders as the most prevalent. Again, *Clostridiales* and *Bacteroidales*, which were among the most abundant orders were composed of low abundant genera. Regarding the standard samples, *Neisseria* was the predominant genus in three of the samples, while *Chitinophagaceae* (*uncl*.) was in another. *Streptococcus*, *Lactobacillus*, *Rothia*, *Moraxella* and *Bergeyella* were among the most abundant genera, with less variability between animals. Although *Clostridiales* and *Bacteroidales* were the 4th and 5th most abundant orders, respectively; they were composed of several low abundant genera.

The differential composition of the microbiota from the deep nose samples was compared with the rest of nasal samples using ANCOM-BC at order, family, genus, species and ASV level. Although the reduced number of samples and the variability between animals from this dataset complicated the search of differently abundant taxa, some differences were identified. Among the differently abundant taxa identified in the external part of the nose, the most relatively abundant were *Flavobacteriales*, *Dyella*, and a highly abundant *Neisseria shayeganii* ASV that was absent in the deep samples. On the contrary, *Pseudomonas* seemed to constitute a bigger portion in the microbiota of the deep nose compared to the other nasal cavities. All the differentially abundant taxa identified comparing the deep and other nasal samples are shown in **Supplementary Table 4**.

#### Supplementary Figures

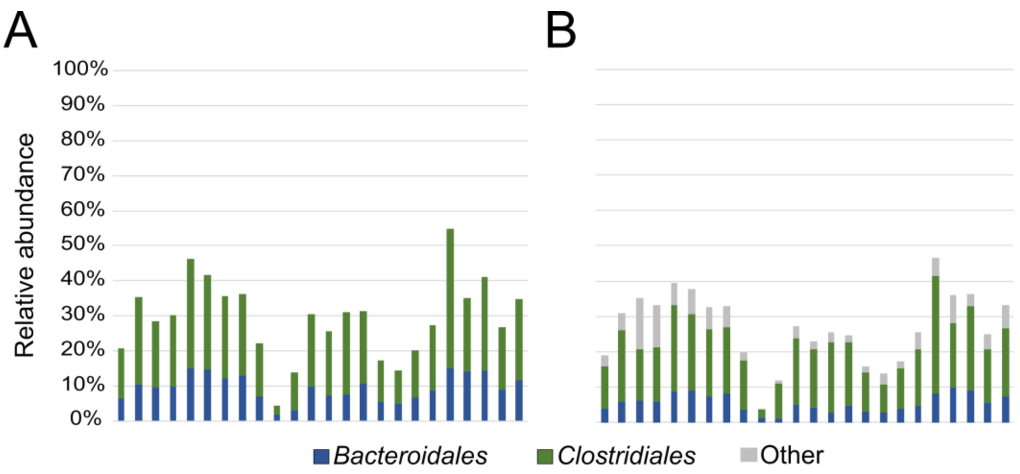


**Supplementary Figure 1. Detection of gut-microbiota associated taxa in nasal swabs. A)** Relative abundance of taxa belonging to *Bacteroidales* (blue) and *Clostridiales* (green) orders in nasal swabs of 24 animals. **B)** Order-level relative abundance of gut-associated taxa using an alternative metric (prevalence of respective family in a reference gut microbiome data set, see **supplementary text 1**). Data are summed by order, namely Bacteroidales (blue), Clostridiales (green), and all other orders (grey).

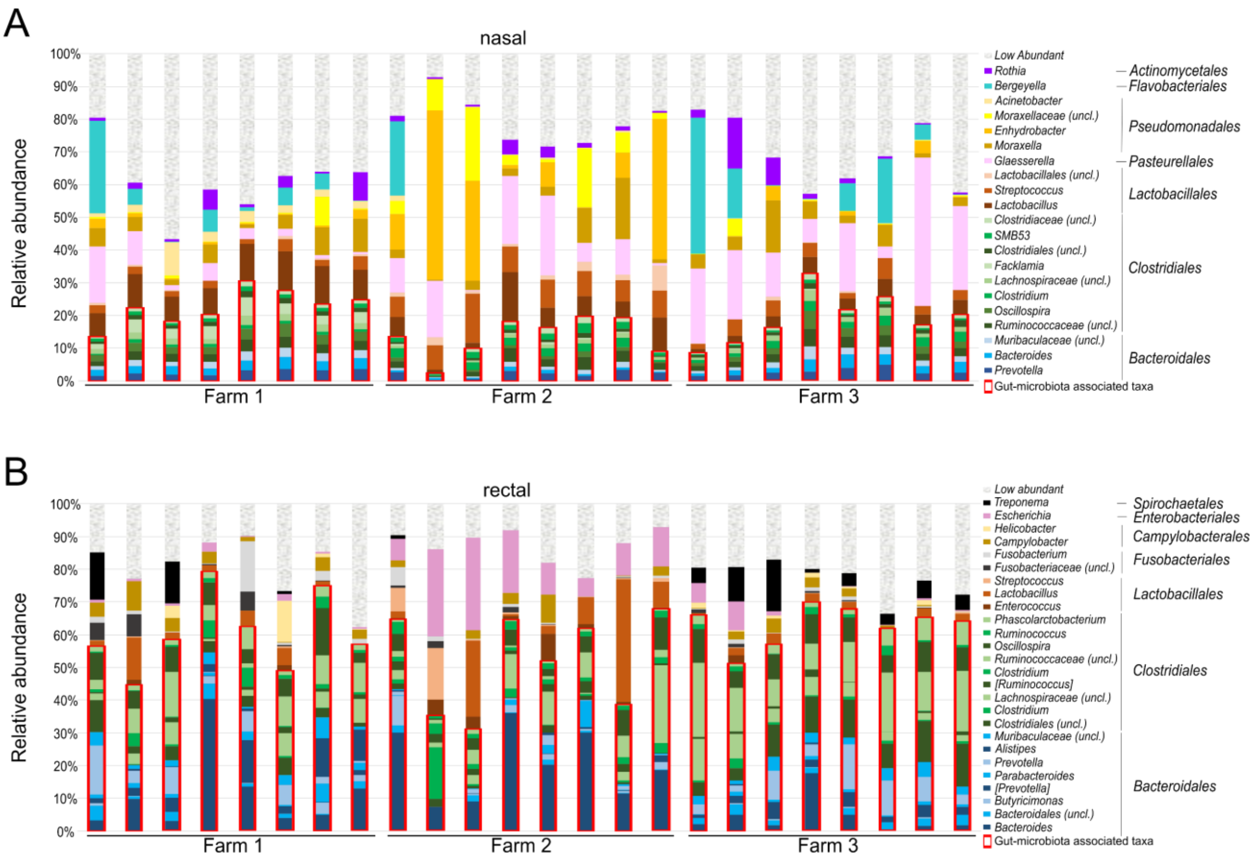


**Supplementary Figure 2. Composition of nasal (A) and rectal (B) microbiota in 24 individual animals across 3 different farms at genus level.** Highlighted in red: gut-microbiota associated taxa. Note that only taxa with > 1% relative abundance are labeled (taxa with <1% relative abundance are summed in the category "Low abundant").

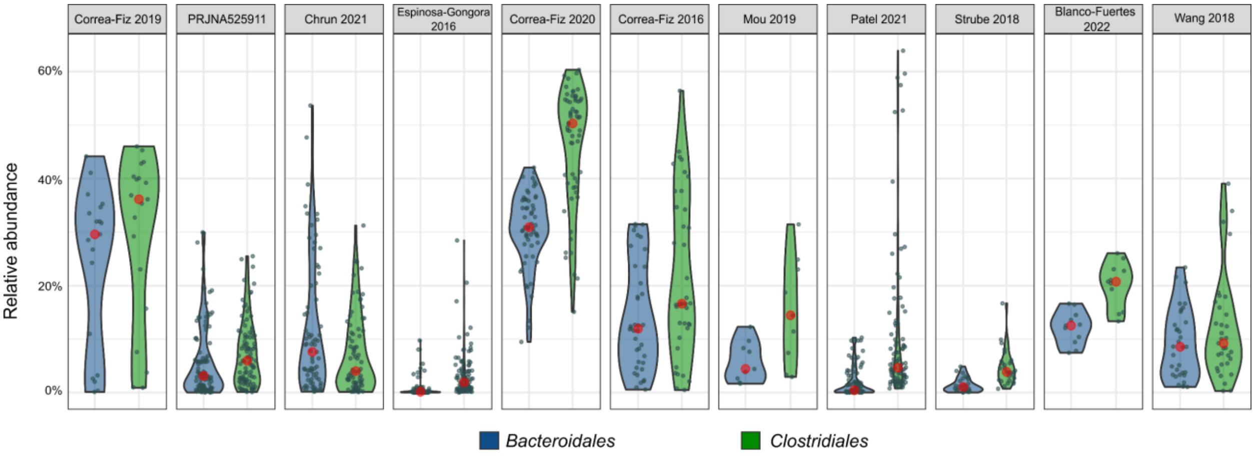


**Supplementary Figure 3. Fraction of gut-microbiota associated taxa (i.e. taxa from *Clostridiales* and *Bacteroidales* orders) in nasal microbiota samples from 11 publicly available data sets**. Data sets are listed in **Supplementary Table 1**. Distribution of data across individual animals is represented in separate violin plots for each data set. Red circle: median. Grey dots: abundance in each individual sample.

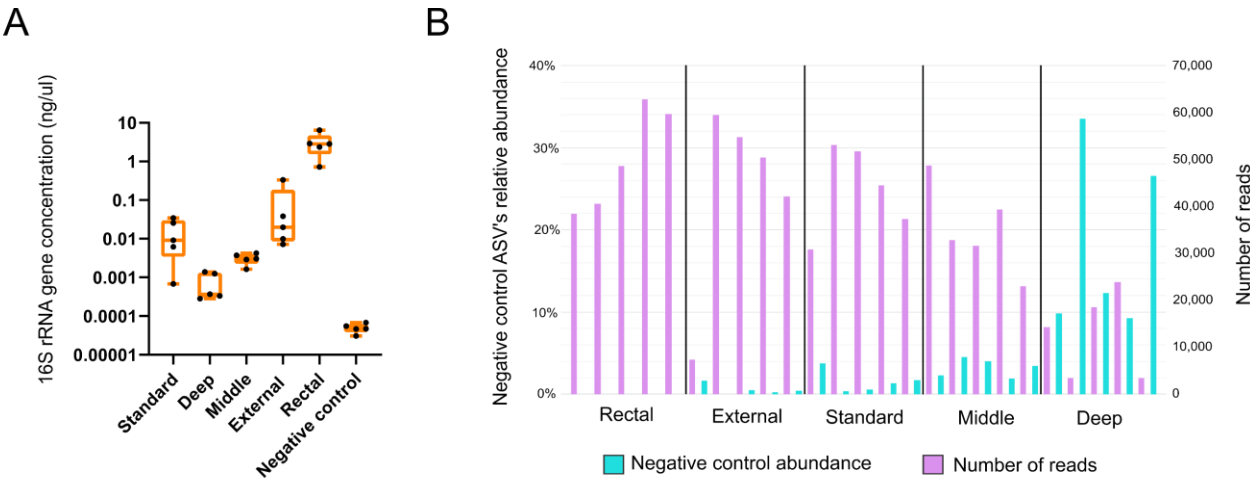


**Supplementary Figure 4. Biomass abundance assessment in surgical samples. A)** Total 16S rRNA quantification of the samples from DNA extracted from the nasal invasive dataset (see methods). In orange, samples obtained from nasal standard swabs. qPCR was performed twice for each sample (data shown is mean of technical replicates). All pairwise differences are statistically significant with Mann-Whitney test ($p$ < 0.05) **B)** Summed relative abundance of the negative control ASVs (turquoise) and number of reads (purple) in each of the same samples.

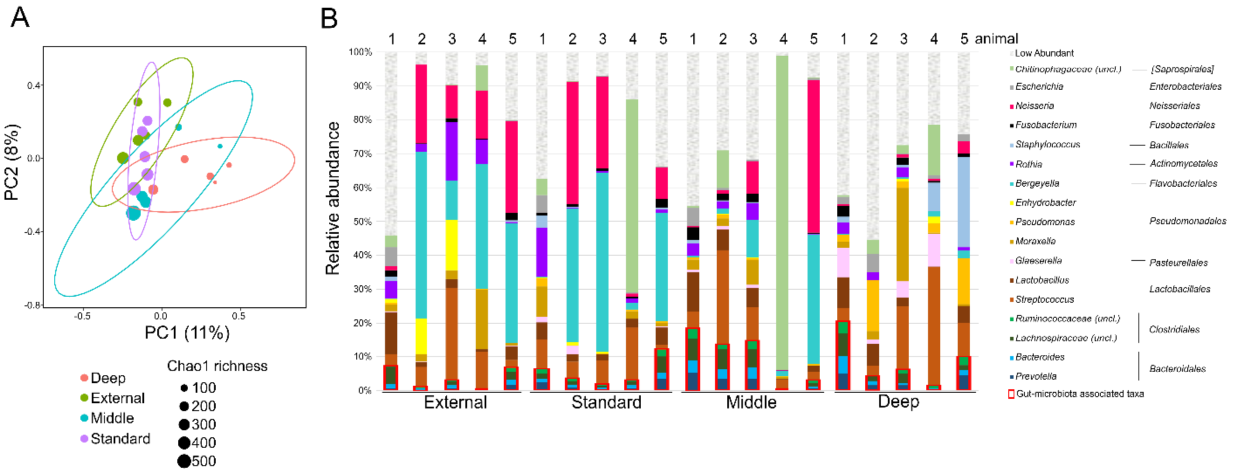


**Supplementary Figure 5. Composition of surgical nasal cavity samples. A)** Principal Component Analysis of microbiota samples performed with Jaccard dissimilarity index (shown here: first two principal components with respective explained variance). Each circle denotes an individual animal, and circle size denotes the alpha diversity estimated by Chao1 richness index. Group ellipses are calculated with the euclidean distances of the samples within each group. **B)** Detailed composition of individual samples at genus level. The order level for each genus is also included in the legend for clarification. Highlighted with red squares: gut-microbiota associated taxa. Note that only taxa with > 1% relative abundance are labeled (taxa with <1% relative abundance are summed in the category "Low Abundant").

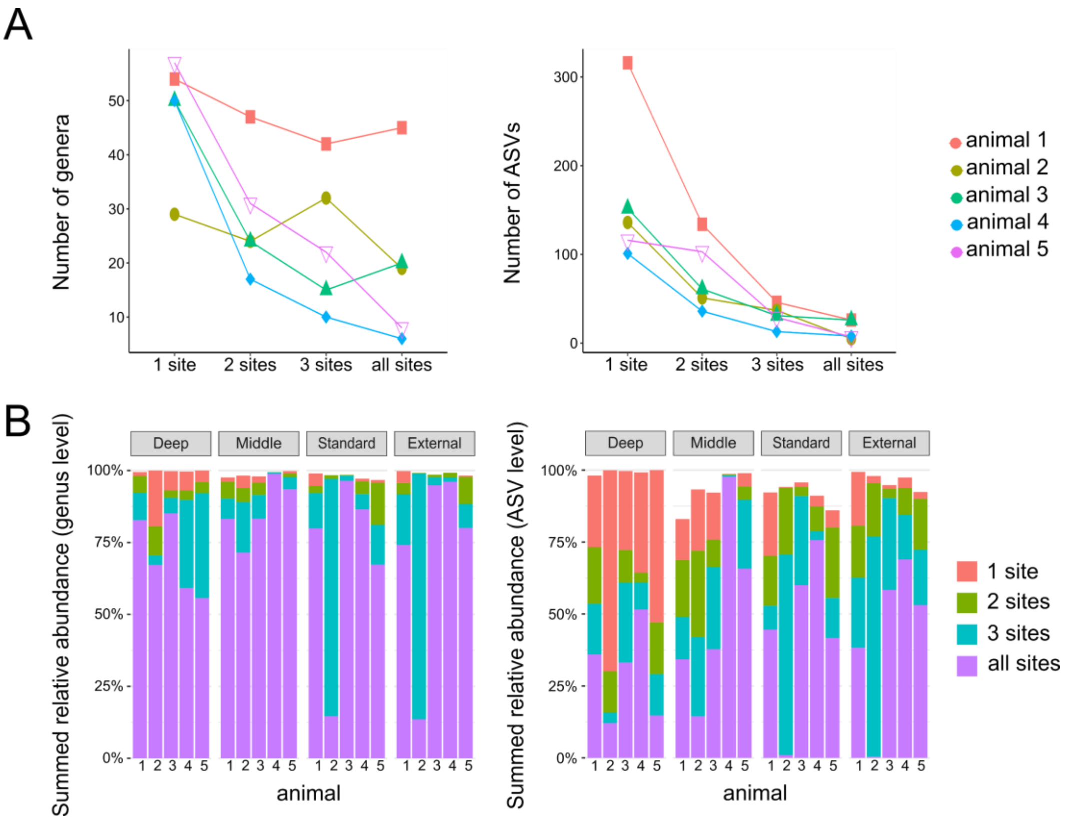


**Supplementary Figure 6. Prevalence of genera and ASVs at different sampling sites. A)** Number of genera (left) or ASVs (right) detected at different sites (from “detected at 1 site only” to “detected at all 4 sites”) in each individual animal by accounting the four types of nasal samples from the invasive dataset (see methods). Only genera and ASVs detected >0.1% relative abundance in at least one sample were considered as “detected”. Genera/ASVs that never exceeded these thresholds are not shown. Note that only a small fraction of genera/ASVs are typically detected at all sites. **B)** Respective summed relative abundance for genera (left) and ASVs (right) detected at different sites (determined as in A). Note that summed relative abundances are dominated by the small fraction of genera/ASVs that are detected at all sites (purple bars).

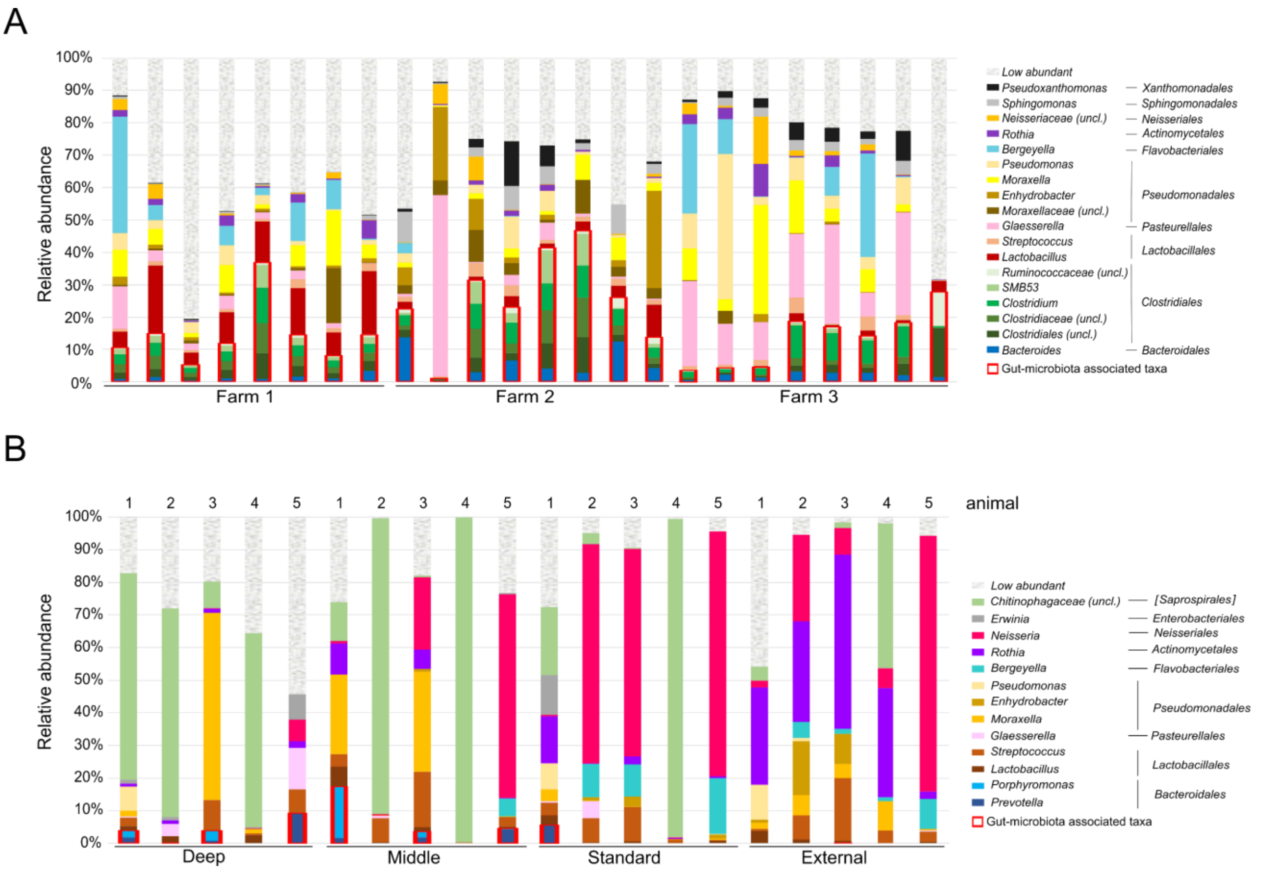


**Supplementary Figure 7. RNA-based Nasal microbiota composition as determined by 16S rRNA cDNA sequencing at genus level. A)** Nasal swabs of 24 animals sampled from 3 farms (see methods). **B)** Surgical samples taken at different sites in the nasal cavity from 5 animals. Highlighted in red: gut-microbiota associated taxa. Note that only taxa with > 1% relative abundance are labeled (taxa with <1% relative abundance are summed in the category "Low abundant").

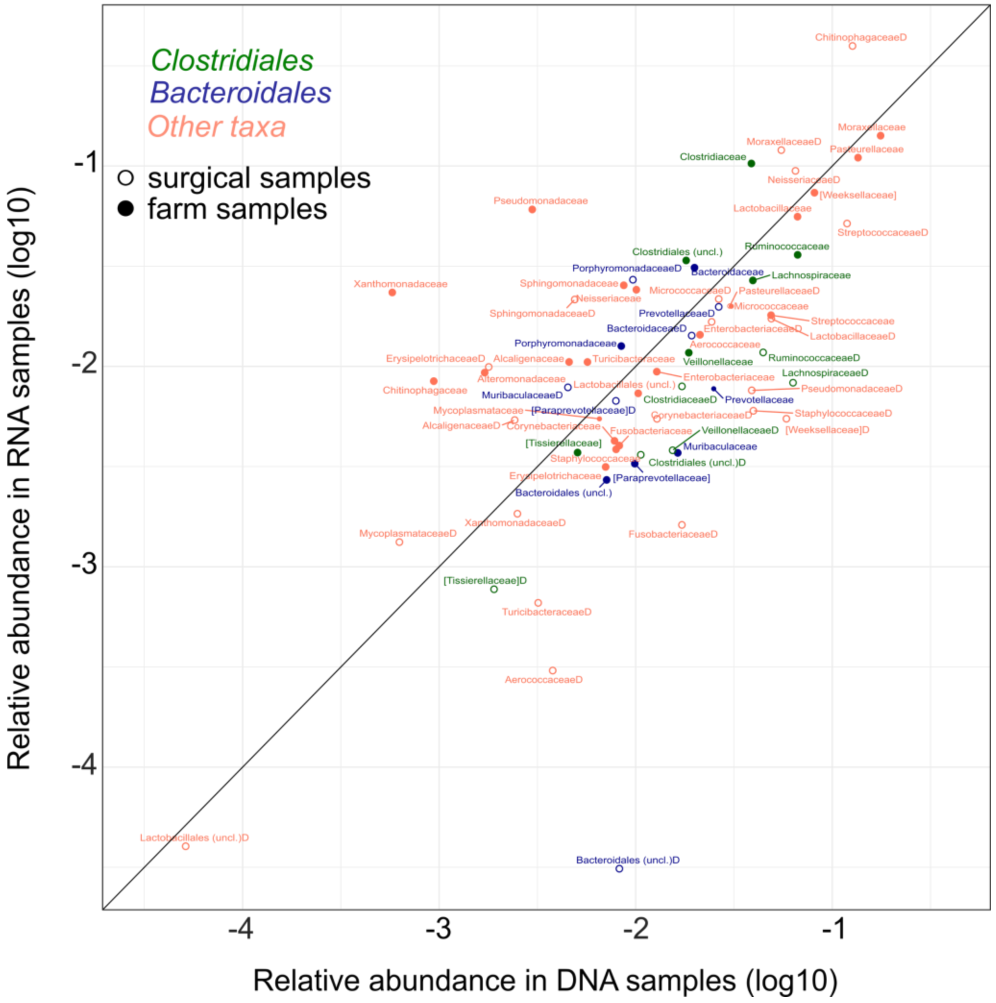


**Supplementary Figure 8. Comparison of 16S rDNA and rRNA mean abundances for the most abundant families in farm and surgical samples.** Relative abundance in DNA and RNA samples was determined by 16S rRNA sequencing of DNA and cDNA, see main text for families with > 0.5% mean abundance in DNA or RNA farm samples. The abundances of the selected families in deep surgical nasal samples are shown as well (with an extra "D" in the label). Gut-microbiota associated taxa (*Bacteroidales* and *Clostridiales*) are labelled in blue and green, respectively. Families from other orders are shown in pink. Open circles: deep nasal samples obtained surgically from 5 animals (deep and middle nasal cavity, see **Figure 3**). Filled circles: standard nasal swabs from 24 animals across 3 farms. Shown are mean values across all samples.

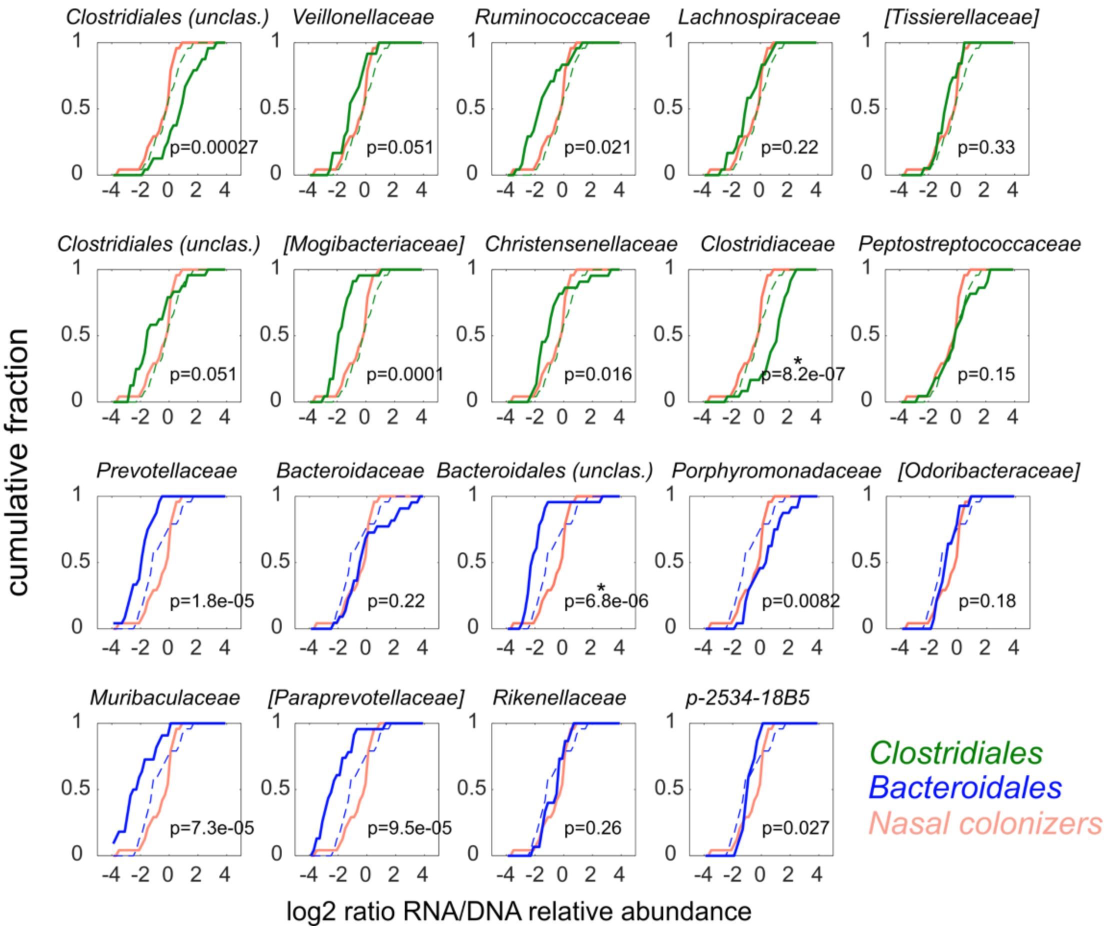


**Supplementary Figure 9. Distribution of RNA/DNA ratios in gut-microbiota associated taxa at family level in nasal swabs from 24 farm animals.** Top two rows: *Clostridiales*, bottom two rows: *Bacteroidales*. Black: ratio for well-established nasal colonizers (combining *Moraxellacae*, *Pasteurellaceae*, *Streptococcaceae, Lactobacillaceae, [Weeksellaceae]*). Green/Blue: ratio for gut-microbiota associated family of interest (dashed line: corresponding order level data). P = p-value of two-sample Kolmogorov-Smirnov test (testing whether the two samples stem from the same continuous distribution). Plots denoted with *: $p < 10^{-5}$. Square brackets in taxonomical assignations indicate contested names in the reference Greengenes database used (Version 13.8). Only taxa which had a relative abundance (at DNA level) > 0.1% in at least 50% of the samples were considered.
